# Behavioral risk models explain locomotor and balance changes when walking at virtual heights

**DOI:** 10.1101/2024.11.18.624196

**Authors:** Nooshin Seddighi, Nicholas J. Woo, Jaylie Montoya, Nicholas Kreter, Mindie Clark, A. Mark Williams, Tiphanie E. Raffegeau, Peter C. Fino

## Abstract

Walking in daily life requires humans to adapt to environments that can influence one’s fear of falling and anxiety about a potential fall. In such environments, individuals may adopt compensatory locomotor and balance changes to maintain a constant expected risk function equal to the product of the probability of some event (e.g., a fall) and the cost of that event (e.g., injury or death). Here, we tested whether locomotor behaviors broadly align with this risk model in two experiments with height-related threats in immersive virtual reality. In Experiment 1, we examined how individuals change their locomotor trajectory while walking along a straight high-elevation walkway. In Experiment 2, we examined how individuals change trajectory and balance control during curved walking where the location of high elevation threat varied. Participants adopted two behaviors that decreased their probability of falling off the edge and aligned with the risk-based model: participants altered their proximity to perceived threats that pose high costs (e.g., a high-elevation ledge), and decreased mediolateral center of mass velocity when that was not possible. Taken together, our results suggest that individuals alter locomotor behavior to change the probability of falling based on the perceived cost of that fall.

## INTRODUCTION

Locomotor control in daily life requires humans to adapt to environmental demands. For instance, changes in terrain alter step length and duration [1], and dynamic obstacle crossings require changes in movement decisions [2]. The effects of external forces or sensory manipulations from the environment are frequently studied in the context of locomotor control [3–9], but environmental features may also elicit compensatory locomotor changes based on psychological factors, including perceived risk and anxiety (e.g., fear of falling when walking along the edge of a cliff) [10]. Theoretical frameworks of fall-related anxiety incorporate experience-driven priors of stability or instability, sensory gains, cortical error responses, and reinvestment-based conscious control to map the effects of perceived threats to balance onto compensatory actions [11, 12]. However, it remains unclear how these threats are systematically evaluated by the person.

Risk-based frameworks can be used to understand human motor control, and similar applications may describe compensatory locomotor changes in response to environmental threats [13]. Classical definitions of risk stem from the maximum expected utility hypothesis, whereby the expected utility or risk of action *a*, *E*[*U|a*] is the product of the probability, *P*(x|a), of some event *x* (e.g., a fall) given action *a* (e.g., a specific foot placement), and the utility or cost of that event, *U*(x), (e.g., injury or death), expressed as *E*[*U*|*a*] = ∑_*x*_ *P*(*x*|*a*) *U*(*x*) [14–17]. The maximum expected-utility framework has been applied in various models of feedback control (e.g., risk-neutral, risk-sensitive), temporal discounting, and diminishing marginal utility to explain motor control of upper limb reaching and static postural tasks [14, 18–21]. Similar probability frameworks may describe compensatory locomotor changes when faced with threats related to falling [21]. For example, humans may adopt locomotor patterns that reduce the probability of falling in environments where the expected cost of a fall (i.e., the likelihood of a severe fall-related injury) is high to maintain a constant expected value of risk.

Compensatory locomotor changes are evident when walking at physically elevated walkways (i.e., real threats) [22–25] or simulated heights in virtual reality (i.e., perceived threats) [26–28]. When walking with height-related threats (i.e., high perceived costs of falling), individuals often exhibit shorter step lengths, wider step widths [29], and changes in the timing of gait phases [30]. Similar changes occur during standing tasks, with individuals adopting “stiffer” control and reduced sway away from the direction of the postural threat (i.e., the edge of the platform) [31, 32]. Such locomotor and postural changes may be an attempt to reduce the probability of falling, *P(x|a)*, in settings where falling is perceived to be particularly harmful (i.e., high *U(x)* where there is a high risk of fall-related injury or death if one falls off of an elevated platform). Under height-related threat, participants often disrupt their execution of familiar, automated movements by unnecessarily “reinvesting” attentional resources toward conscious movement regulation [33]. Because gait involves relatively little conscious control in healthy populations, anxiety-induced reinvestment is seen as unnecessary and often leads to behaviors indicative of cautious gait [29, 34, 35]. Despite prior work on standing and walking at virtually or physically elevated heights, it remains unclear whether such locomotor changes align with risk-based models. Few, if any, studies have characterized measures that directly indicate the probability of falling to identify risk-based locomotor behavior.

The purpose of this study was to test whether individuals’ behavior aligns with a risk-based model of locomotor control when walking in a virtual environment with simulated height-related threats. We hypothesized that, when presented with a height-related threat (a ∼15 m virtual cliff, in this case), individuals would exhibit locomotor changes that decrease the probability of falling off the walkway toward the cliff (i.e., decrease *P(x|a)* in the direction with greatest perceived cost *U(x)*; Figure 1). We tested this hypothesis using two experiments with virtual height-related threats. In Experiment 1, we examined how individuals change their locomotor trajectory at virtual high-elevation to regulate risk during a straight locomotor task. In Experiment 2, we examined how individuals change both locomotor trajectory and balance control at virtual high-elevation to minimize risk during a turning locomotor task. We predicted that participants would adopt compensatory trajectories (e.g., increase the distance between themselves and the threatening side of the walkway) and adjust balance control (e.g., decrease the probability of falling towards the threatening side of the walkway) to minimize the probability of falling in the direction of the greatest threat.

**Figure 1.**
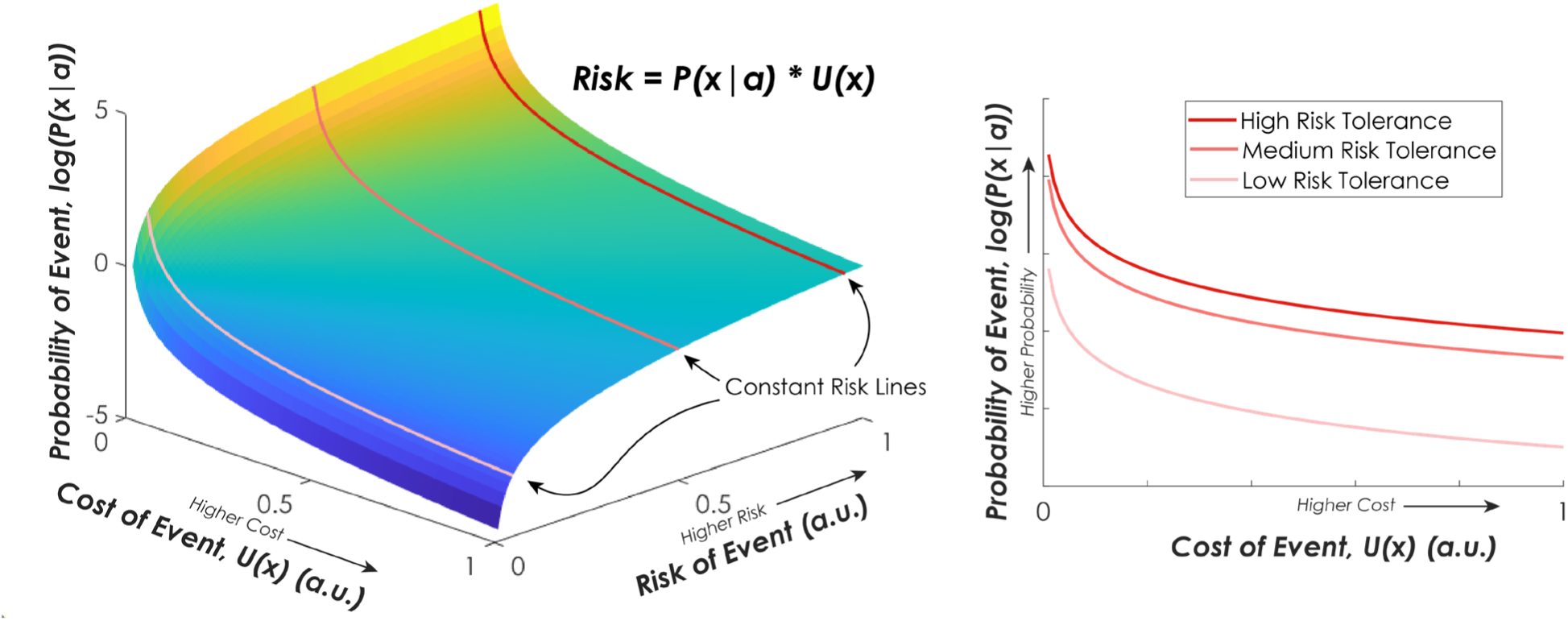
Conceptual model of a risk surface for a single action *a* based on *E*[*U*|*a*] = ∑_*x*_ *P*(*x*|*a*) *U*(*x*) where the risk is based on the expected value *E*[*U*|*a*] equal to the product of the probability of an event given that action, *P*(*x*|*a*) and the cost, or consequence, of that event, *U*(*x*). If decisions are based on regulating risk, then lines of constant risk may indicate an individual’s risk tolerance, and behavioral modifications could select actions based on the perceived cost of an event (i.e., fall) given some action. For example, individuals compensate for events with high expected cost (e.g., falls from high heights) by selecting actions that have a low the probability of those high-cost events occurring.

## METHODS

### Participants

Two different groups of participants were recruited for Experiments 1 and 2. All participants provided informed written consent in compliance with the University of Utah’s Institutional Review Board. The inclusion criteria for both experiments were (1) age 18 – 65 years old, (2) ability to walk without aid or discomfort, (3) capacity to follow verbal commands, (4) could comfortably wear the VR-head mounted display (HMD), (5) had normal or corrected to normal hearing and vision. The exclusion criteria were (1) a history of neurological disorders that may impair movement, (2) diagnosed non-medically stable cardiovascular disease, (3) recent orthopedic injury or surgeries in the last year, (4) chronic dizziness or excessive motion sickness, (5) pregnant women for balance considerations.

### Virtual Environment

Participants completed a series of walking tasks in an immersive virtual reality (VR) environment to elicit changes in their perceived height-related threat of falling. Participants wore their normal corrective eyewear and were fitted with a wireless HTC Vive (version 2.0, Bellevue, WA) HMD that displayed a virtual walkway that matched the physical walkway in the laboratory. Two VR trackers (HTC VIVE Tracker 2.0) were placed around the lateral ankle, proximal to the lateral malleolus. Virtual representations of the feet (tennis shoe avatars) were provided to the participant using these ankle-mounted trackers following a calibration procedure.

### Experiment 1

#### Procedures

Experiment 1 used a rectangular walkway that was 1.22 m wide, 2.44 m long, and 1.90 cm high (Figure 2). The virtual walkway was matched to a physical walkway (height: 1.90 cm) in the laboratory to provide proprioceptive feedback of the walkway edge. During the walking task, two different virtual conditions were presented to the participants in a randomized order: a high walkway (15 meters in the air) and a low walkway (ground level) height. In the Low height condition, the plank was virtually presented at ground level. In the High height condition, the plank was virtually elevated 15 m above the ground.

**Figure 2.**
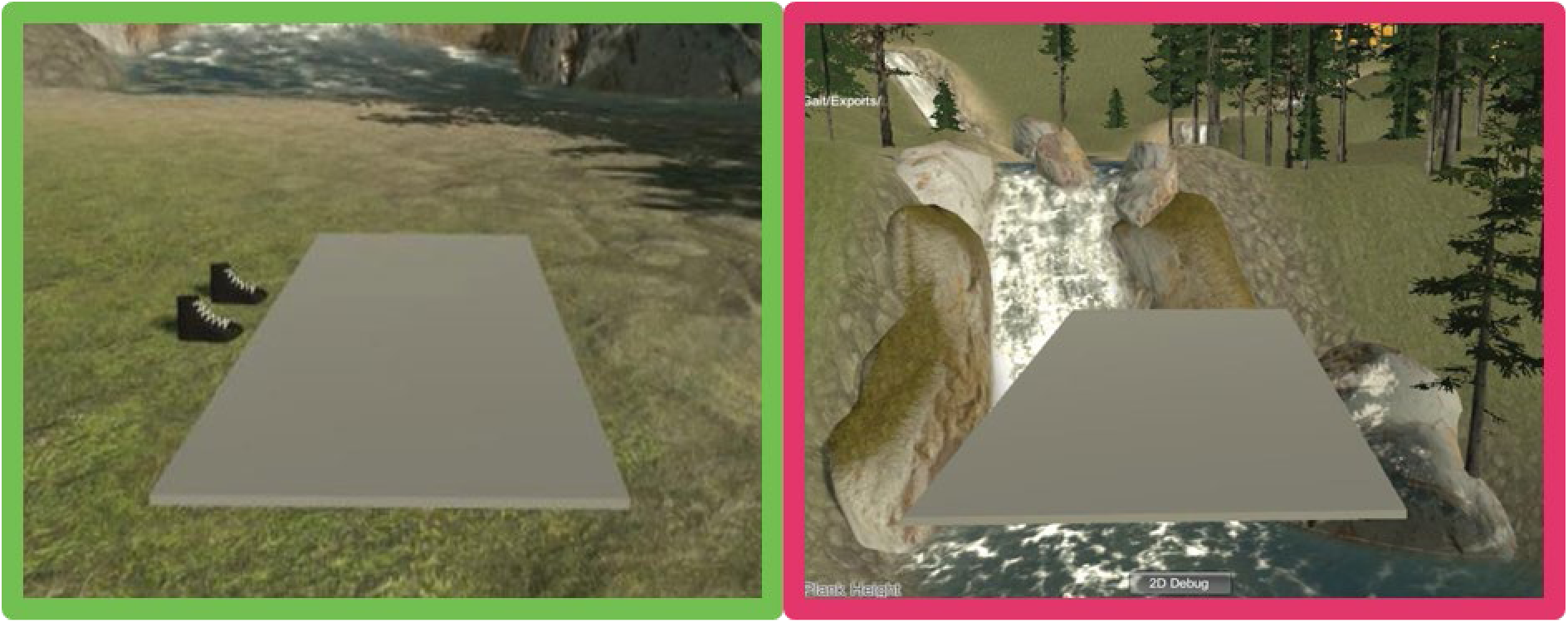
Experimental setup for Experiment 1 depicting the Low (left) and High (right) height conditions within the VR environment.

To standardize the starting location for each trial, participants were asked to begin each walking trial by facing the length of the walkway and standing at the corner such that their outer foot’s heel aligned with the corner of the walkway and the lateral edge of their foot ran along the outer edge of the physical walkway. Participants began each condition standing at the designated starting position; the walkway was then elevated to the desired virtual height (15 m at a 1 m/s rate). Participants were then instructed to walk to the opposite end of the walkway and come to a stop when they reached the end. They were instructed not to adjust their foot placement once they reached the end of the walkway. Once participants came to a stop, their final foot placement was recorded on the walkway. Participants were then asked to turn around, place their other foot in the standard starting position aligned with the corner of the plank on the same edge they previously started, and the process was repeated. Each virtual height condition started with the participant’s outer edge of their foot aligned with the edge of the walkway. Conditions contained two trials – walking down and back on the walkway – with either the left or right foot aligned with the edge of the walkway. After completion of the two trials within each condition (down and back along the walkway), the virtual walkway was lowered to the ground (15 m at a −1 m/s rate).

Threat conditions were presented in three blocks, each containing 8 total trials (2 virtual height conditions x 2 initial starting positions x 2 trials per condition). The order of the conditions within the first block was kept constant to ensure the Low condition was always presented first, but the initial starting position (i.e., starting with either the right or left foot aligned with the walkway edge) was randomized. In the two remaining blocks, height conditions and initial starting positions were randomized, such that either High or Low conditions could come first. A total of three blocks with four conditions were collected for each participant, yielding 24 walking trials (12 per side) per participant.

#### Primary Outcomes

The primary outcome measure for Experiment 1 was lateral deviation, or movement along the width of the walkway. Lateral deviation represents compensatory, unnecessary motion that is orthogonal to the explicit goal of walking the length of the walkway and away from the optimal shorter path to the desired destination. The lateral deviation was measured from the outer edge of the physical walkway to the outer foot’s lateral edge after each trial.

### Experiment 2

#### Procedures

Experiment 2 used a virtual circular ring walkway with a 0.4 m width (1.2 m inner-radius and 1.6 outer-radius). The virtual walkway had a matching 1.27 cm high physical walkway in the laboratory to enable proprioceptive feedback of the walkway edge (Figure 3a). In addition to the two VR trackers placed laterally around the ankles, two VR hand controllers (HTC Vive) were placed in a waistbelt and located over the approximate location of navel and L2-L3 lumbar vertebrae to capture estimated center-of-mass (CoM) kinematics using the average of the two VR hand controllers’ position.

**Figure 3.**
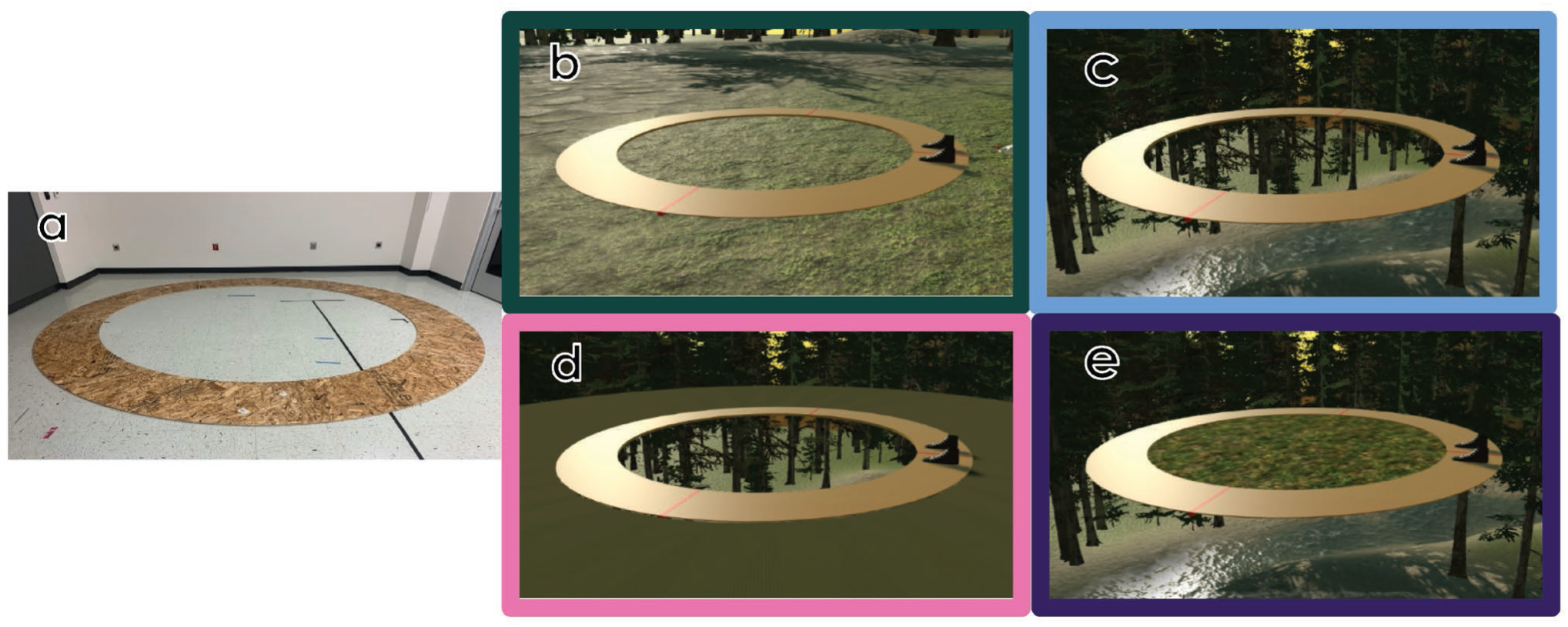
Virtual and physical circular walkways for Experiment 2 depicting the circular walkway in the physical world (a) and four immersive environments: (b) ground level (No Threat), (c) high elevation with both threats present (Bilateral Threat), (d) high elevation with the inner threat present (Inner Threat), and (e) high elevation with the outer threat present (Outer Threat).

Participants were given a brief acclimation period to adjust to the VR environment, which included a procedure to obtain their self-selected walking speed in the low-elevation VR environment. For this speed-calibration procedure, participants were instructed to walk four laps around the circular walkway at their comfortable, self-selected walking pace. For all walking trials, participants walked in a direction such that their dominant leg was on the outside of the circle (i.e. if the participant was right foot dominant, they walked in a counter-clockwise direction). After a brief (<1 minute) rest, participants were asked to repeat the task two more times, resulting in three speed-calibration trials. The total duration of the middle two laps in each condition was recorded and averaged across the trials to generate a fixed self-selected walking speed for each participant. To ensure walking speed remained constant across all remaining trials, participants were paced by a metronome that was delivered at ¼ lap intervals based on their self-selected speed described above. Participants were instructed to traverse ¼ of the walkway (marked with red lines at ¼ intervals) for each metronome cue [36, 37].

Participants completed four 40-second walking trials within the VR environment at their constant, self-selected speed: (1) walking at ground level (No Threat - **Figure 3b**); (2) walking at high elevation with a threat on both sides (Bilateral Threat - **Figure 3c**); (3) walking at high elevation with an inner threat (Inner Threat - **Figure 3d**); and (4) walking at high elevation with an outer threat (Outer Threat - **Figure 3e**). The Bilateral, Inner, and Outer threat conditions presented the circular walkway suspended approximately 15 meters above the ground. The No Threat condition was always presented first; the order of the remaining three conditions (Bilateral, Inner, Outer) was randomized across participants. The 40-second duration was selected to limit patient-reported dizziness.

Participants began each condition seated on a chair, placed on the walkway, facing the tangential direction of the walkway, and viewing the No Threat condition. For the Bilateral Threat, Inner Threat, and Outer Threat conditions, the participant was then virtually elevated to the desired height (∼15 m at a 1 m/s rate). After the desired height was reached, participants were asked to stand, and a research assistant removed the chair from the walkway. Participants were instructed to walk around the walkway and match the pace of the metronome from the speed-calibration for 40 seconds. At the completion of each trial, a research assistant replaced the chair, the participant returned to a seated position, and the virtual walkway was lowered to the ground (15 m at a −1 m/s rate).

#### Data Analysis

The motion trackers recorded positional data using four lighthouse-based infrared sensors at an average sampling frequency of 70.58 Hz, where the center of the walkway served as the origin of the positional data. The raw positional data were extracted for each tracker and resampled at a constant 200 Hz using the ‘resample’ command in MATLAB with linear interpolation. The resampled data were then filtered using a 6 Hz low-pass, phaseless 4^th^ order Butterworth filter. The three-dimensional position vectors of the two body trackers were averaged to yield approximate position and velocity vectors of the CoM, where velocity was calculated using the central difference method [38].

Heel contact and toe-off gait events were defined using the difference in the transverse plane between the foot and CoM, corrected for the turning gait [39, 40]. To account for changes in heading angle that modify the coordinate system, a time-varying, walkway-fixed local reference frame was defined using the radial vector from the center of the circular walkway to the position of the CoM, a vertical axis, and the right-handed cross-product of the two [37]. Using a walkway-fixed reference frame defined the anteroposterior (AP) and mediolateral (ML) directions for decomposing position and velocity vectors.

#### Primary Outcomes

Since our focus was on the threat of falling off the edge of the walkway, the radial distance from the edge of the walkway to the extrapolated CoM (XcoM) at contralateral toe-off – hereafter referred to as the margin of falling (MoF) was calculated as the primary outcome variables for each condition. The XcoM was calculated using the position and velocity of the 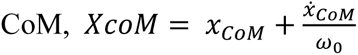, where ω_0_ is the eigenfrequency calculated using 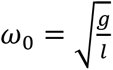, where *g* is the gravitational constant, and *l* was the pendulum length, defined here has the distance from the estimated position of the CoM to the foot tracker on the lateral malleolus [41, 42]. This MoF, calculated as the radial difference between the edge of the walkway and the XcoM is conceptually similar to the margin of stability, which quantifies the instantaneous mechanical stability of an inverted pendulum [43]. In this context, a larger positive MoF indicates the XcoM is positioned further from the edge of the walkway in question. If the margin of stability was negative, a positive MoF would represent a larger distance between the XcoM and the edge of the walkway, enabling cross-over steps to prevent oneself from falling off the edge. A negative MoF indicates a smaller distance between the XcoM and the edge of the walkway, such that the participant would fall off the edge unless stance-phase counter-rotation strategies or other external forces are applied. The MoF was extracted for each step throughout the trial at contralateral toe-off. To eliminate transient effects associated with the start and stop of the trial and ensure participants were matching the metronome cadence, the first 8 steps on each limb and the last step on each limb were excluded from analysis.

To directly quantify the probability of falling off the edge of the walkway (PoF), we adapted the probability of instability (PoI) originally described for margin of stability [44]. To calculate the PoF, we integrated the probability density function of the MoF for steps in each trial and computed the likelihood of falling off the edge of the walkway (without counter-rotation measures) for any given step using the cumulative probability of the MoF being less than 0. Our approach is analogous to the original PoI definition [44], but here it directly quantifies the probability of falling off the edge of the walkway rather than the probability of mechanical instability that necessitates a corrective action.

#### Secondary Outcomes

We measured the average radial distance of each step from the center of the circle for every step in the trial and the mediolateral (ML) CoM velocity, measured at contralateral toe-off, as secondary outcomes for Experiment 2. The radial distance of each step was calculated as the distance from the center of the circular walkway to the position of the foot trackers during each stance phase. The ML CoM velocity was defined using the walkway-fixed reference frame, making it synonymous with the radial CoM velocity [37].

### Statistical Analysis

To confirm the effectiveness of our virtual height manipulation at eliciting height-related anxiety, the Mental Readiness Form-3 (MRF-3) and Rating Scale of Mental Effort (RSME) were administered [45, 46]. In Experiment 1, the MRF-3 and RSME were administered after every walking pass in the first and third trial blocks (i.e., after every walk from one end of the walkway to the other). In Experiment 2, the MRF-3 and RSME were administered after every condition (i.e., after every 40 s walking trial; see Supplement A). Participants remained in the virtual environment while completing these questionnaires; the MRF-3 was administered verbally and the RSME was displayed in the virtual environment to ensure participants had accurate anchors for their responses. We report the findings from the MRF-3 and the RSME in the supplemental material (see Supplements A).

#### Experiment 1

To test whether individuals altered their locomotor trajectory away from the edge of the walkway when there was a height-related threat, the average lateral deviation for Low and High conditions was computed, condensed across side of threat (left vs. right), and compared using paired sample t-tests. Within-subject effect sizes for the difference between Low and High threat conditions were estimated using Hedges’s *g* [47].

#### Experiment 2

To test whether participants exhibited different balance control when walking with height-related threats, we fit linear mixed-effects regression models (LME) with main effects of condition (No Threat, Bilateral Threat, Outer Threat, Inner Threat), with the No Threat condition treated as the reference condition. Models were adjusted for age and body mass index (BMI) and stratified by stance limb (inner limb vs. outer limb). Models included random intercepts by subject to account for within-subject correlations. Post-hoc pair-wise contrasts were implemented to test for differences between No Threat and Bilateral Threat conditions (to determine the effect of walking at height), and between the Inner and Outer Threat conditions (to determine the effect of the location of the threat). Due to skewness, raw PoF values were log-transformed prior to statistical analysis, with zeros replaced by the minimum non-zero value. Hedges’s *g* effect sizes were computed for all pair-wise comparisons. All statistical analyses were completed in MATLAB R2024a (The Mathworks, Inc. Natick MA, USA).

## RESULTS

### Experiment 1

Six healthy, young adult participants participated in Experiment 1 [4F / 2M; mean (SD) age = 21.5 (0.8) years; height = 1.78 (0.54) m; mass = 76.4 (15.9) kg)]. All participants increased the lateral deviation from the edge of the walkway when walking in the High condition compared to the Low condition (Figure 4; Low mean (SD) = 11.00 (2.47) cm; High mean (SD) = 13.86 (4.25) cm; p = 0.027, Hedge’s *g* = 0.923). Participants reported feeling more worried, tense, and less confident in the high height condition, which required higher mental effort to complete the tasks of Experiment 1 (see MRF-3 and RSME results in Supplement A).

**Figure 4.**
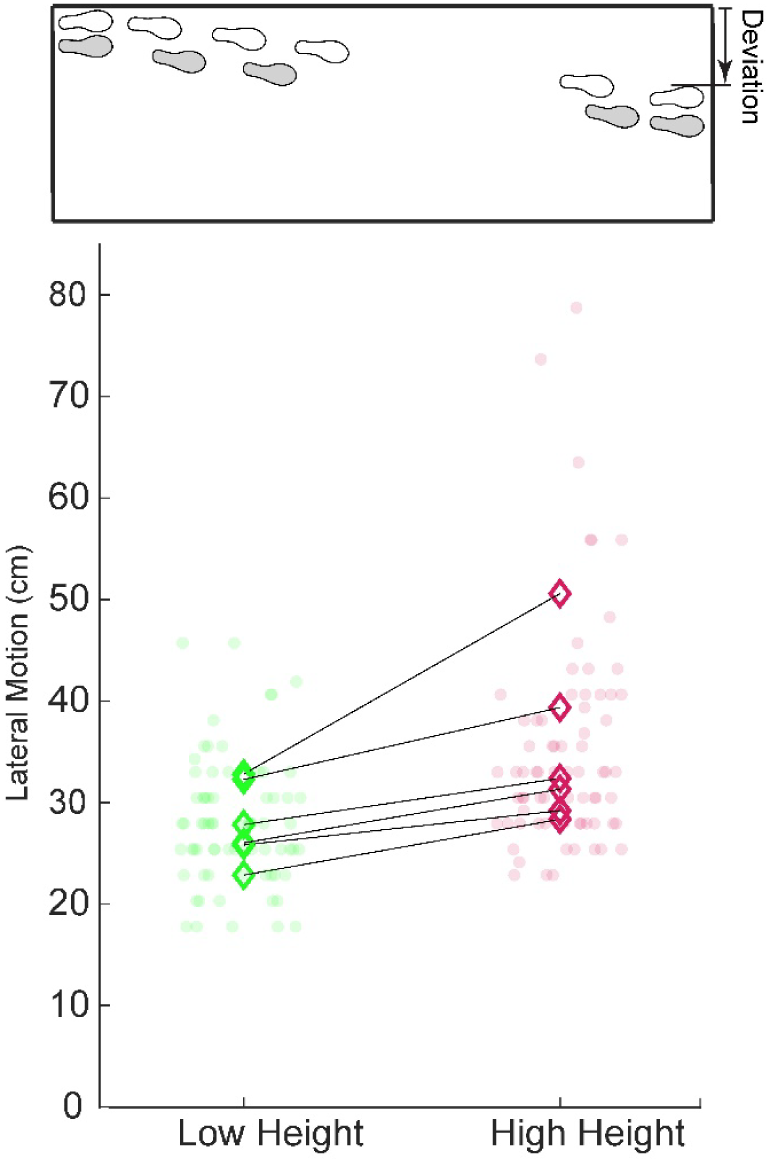
Lateral deviation from the edge across different platform heights in Experiment 1. The top schematic illustrates the initial and final foot positions along the walkway, with arrows indicating the deviation trajectory. The bottom scatter plot shows the lateral motion (in cm) for each subject under low height (green) and high height (red) conditions. Data points represent individual measurements, and the connected diamonds indicate within-participant mean values for each condition.

### Experiment 2

A total of 15 (7 males / 8 females) participants were recruited [mean (SD) age = 25.07 (2.66) years; height = 1.73 (0.08) m, weight = 73.07 (16.12) kg]. Out of these participants, 8 reported experiencing visual height intolerance prior to the experiment. All participants reported right foot dominance. Participants’ mean (SD) self-selected walking speed across each trial was 0.89 (0.14) m/s (No Threat), 0.85 (0.12) m/s (Bilateral Threat), 0.85 (0.12) m/s (Inner Threat), and 0.87 (0.12) m/s (Outer Threat). Additionally, the average (SD) number of steps per condition was 54.3 (7.7) (No Threat), 57.4 (7.6) (Bilateral Threat), 55.3 (8.2) (Inner Threat), and 55.1 (9.2) (Outer Threat).

#### Margin of Falling

Participants’ MoF was significantly influenced by the presence and direction of height-related threats (Table 1, Figure 5). When exposed to bilateral threats, participants decreased their MoF toward the inner edge (*β* = 0.017, SE = 0.006, p = 0.006) and they increased their MoF toward the outer edge (*β* = 0.025, SE = 0.006, p < 0.001) compared to the ground-level condition. When exposed to unilateral threats, participants increased their MoF in the direction of the threat (*β* = 0.062, SE = 0.006, p < 0.001 for the inner limb; *β* = 0.063, SE = 0.006, p < 0.001 for the outer limb).

**Figure 5.**
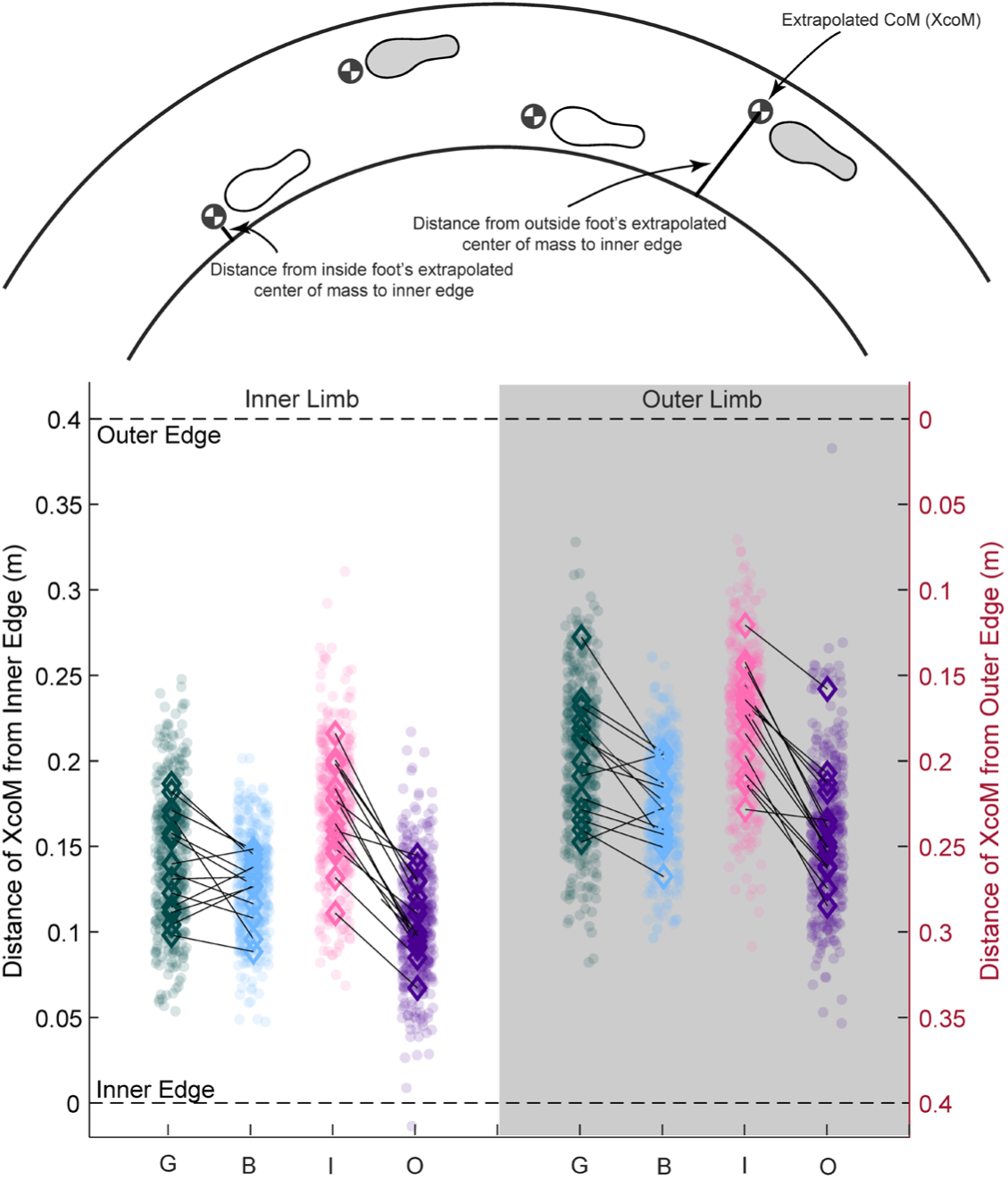
Margin of Falling (MoF) results for Experiment 2 representing the distance of extrapolated center of mass (XcoM) from the inner (white shading) and outer (gray shading) stance limb (i.e., the stance limb closest to the inner and outer edge, respectively) at the time of contralateral toe-off. Conditions include no threat (G), bilateral threats (B), inner threat (I), and outer threat (O). Data points represent individual MoF measures for each step across all participants, and the connected diamonds indicate within-participant mean values for each condition.

**Table 1.** Descriptive means (M) and standard deviations (SD), along with adjusted p-values and effect sizes for each variable, margin of falling (MoF), probability of falling (PoF), radial foot placement, and mediolateral (ML) center of mass (CoM) velocity, across different threat conditions and stance limb. Bolded p-values and effect sizes indicate statistically significant differences.

|  | Stance Limb / Edge | Ground Level M (SD) | Bilateral Threat M (SD) | *Pairwise Contrast P-value* | Hedge's g ES | Inner Threat M (SD) | Outer Threat M (SD) | *Pairwise Contrast P-value* | Hedge's g ES |
| --- | --- | --- | --- | --- | --- | --- | --- | --- | --- |
| Margin of Falling (MoF) (m) | Inner | 0.14 (0.03) | 0.13 (0.02) | **0.006** | **0.67** | 0.17 (0.03) | 0.11 (0.02) | **<0.001** | **2.35** |
|  | Outer | 0.20 (0.03) | 0.22 (0.02) | **<0.001** | **0.84** | 0.18 (0.03) | 0.24 (0.03) | **<0.001** | **1.97** |
| Probability of Falling (PoF) (%) | Inner | 4.62e-03 (1.21e-02) | 1.19e-04 (3.35e-04) | **0.034** | **0.67** | 2.44e-04 (8.31e-04) | 6.34 e-02 (1.22e-01) | **<0.001** | **0.98** |
|  | Outer | 7.62e-06 (2.64e-05) | 4.59e-14 (1.47e-13) | **<0.001** | **2.31** | 7.77e-05 (2.17e-04) | 5.02e-04 (1.59e-03) | **0.002** | **0.89** |
| Radial Foot Placement (m) | Inner | 1.36 (0.04) | 1.36 (0.03) | 0.275 | 0.22 | 1.39 (0.04) | 1.33 (0.03) | **<0.001** | **1.71** |
|  | Outer | 1.45 (0.03) | 1.42 (0.03) | **<0.001** | **0.96** | 1.47 (0.04) | 1.40 (0.04) | **<0.001** | **1.75** |
| ML CoM Velocity (m/s) | Inner | 0.12 (0.05) | 0.10 (0.04) | **<0.001** | **0.38** | 0.11 (0.05) | 0.11 (0.05) | 0.281 | 0.10 |
|  | Outer | 0.10 (0.04) | 0.09 (0.04) | 0.090 | 0.22 | 0.10 (0.05) | 0.10 (0.05) | 0.938 | 0.01 |

Margin of Falling (MoF): Represents the radial distance from the edge of the walkway to the extrapolated Center of Mass (XcoM) at contralateral toe-off, indicating the stability margin in each condition.
Probability of Falling (PoF): The estimated probability, expressed as a percentage, that a participant will fall under each condition.
Medial-lateral velocity of the Center of Mass (ML CoM Velocity): The medial-lateral velocity of the Center of Mass (CoM), measured in meters per second (m/s).

#### Probability of Falling

Participants exhibited changes in their PoF that were aligned with the hypothesized relationship, where the PoF was lower the perceived consequence of a fall was greater (i.e., under threat conditions; Figure 6, Table 1). In the bilateral threat scenario, participants reduced their PoF in both directions compared to the ground-level condition (*β* = 0.005, SE = 0.021, p < 0.034 for the inner limb; *β* = 7.62e-06, SE = 2.76e-04, p < 0.001 for the outer limb). In the presence of unilateral threats, participants exhibited a lower PoF in the direction of the threat for both the inner (*β* = 0.063, SE = 0.021, p < 0.001) and outer limbs (*β* = 4.25e-04, SE = 2.76e-04, p = 0.002), and a higher PoF in the direction opposite of the threat.

**Figure 6.**
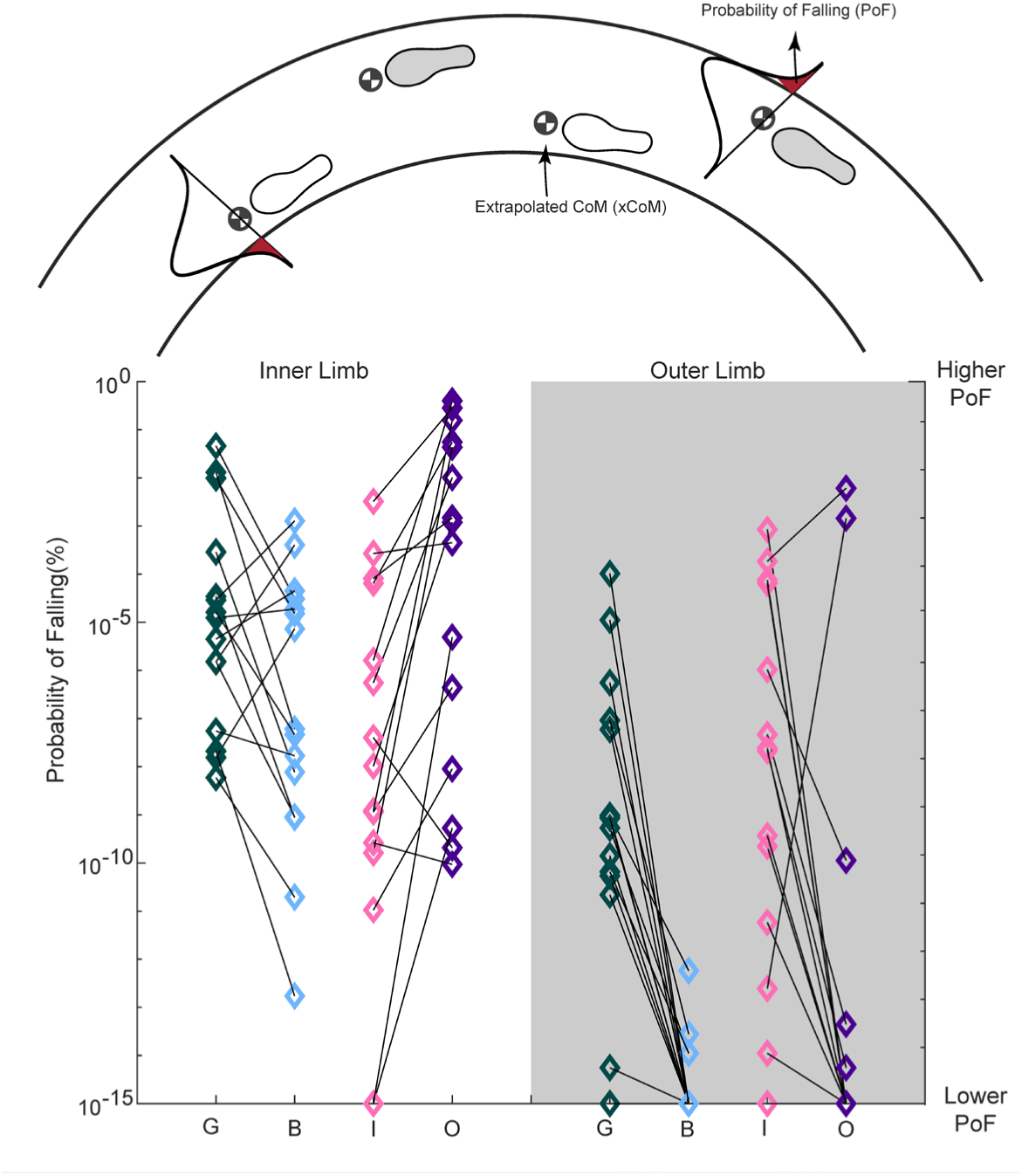
Probability of Falling (PoF) results for Experiment 2 for ground-level (G), bilateral threats (B), inner threat (I), and outer threat (O) conditions for the inner (white shading) and outer limb (gray shading). Data points represent individual MoF measures for each step across all participants, and the connected diamonds indicate within-participant mean values for each condition. Larger values indicate a greater probability of falling off the corresponding edge of the walkway.

#### Radial Foot Placement

The radial placement of participants’ feet, indicating their position on the walkway and step width, was affected by bilateral and unilateral threats differently (Figure 7). When comparing the bilateral threat condition to the ground-level condition, participants’ outer limb was located close to the center of the walkway in the bilateral threat condition (i.e., further from the outer edge) (*β* = 0.031, SE = 0.006, p < 0.001); there was no change in the position of the inner foot (*β* = 0.007, SE = 0.007, p = 0.275). Under unilateral threat conditions, participants shifted both their inner and outer foot placement away from the threat (i.e., moving away from the inner edge of the walkway during inner threats, and moving towards the inner edge of the walkway during outer threats) (*β* = 0.059, SE = 0.007, p < 0.001 for the inner limb; *β* = 0.066, SE = 0.006, p < 0.001 for the outer limb).

**Figure 7.**
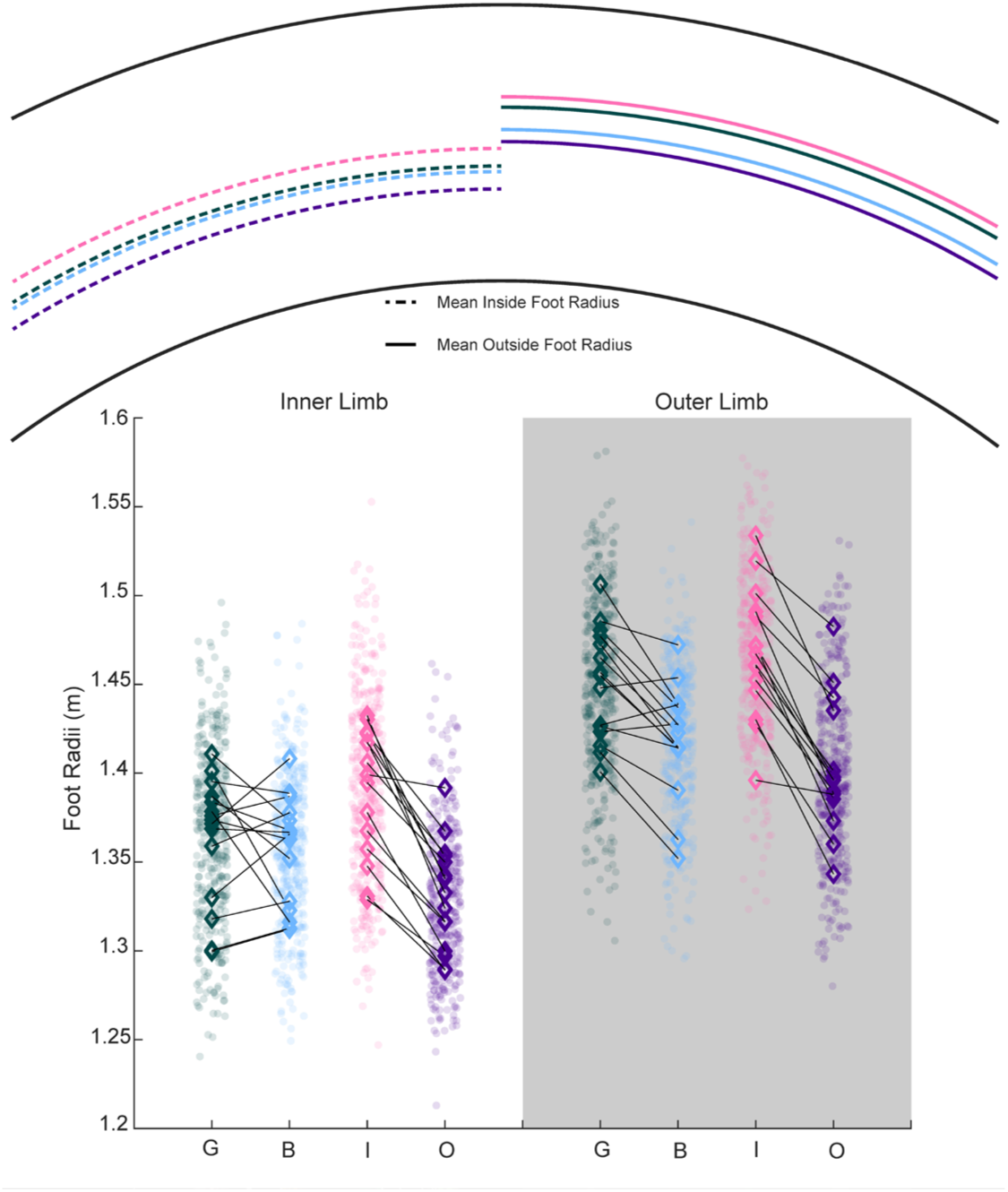
Radial foot placement results for Experiment 2 during ground-level (G), bilateral threats (B), inner threat (I), and outer threat (O) for the inner (white shading) and outer limb (gray shading). Data points represent individual MoF measures for each step across all participants, and the connected diamonds indicate within-participant mean values for each condition. The bounds of the walkway were 1.2 m (inner edge) and 1.6 m (outer edge), with 1.4 m indicating the center of the walkway.

#### Mediolateral Velocity of Center of Mass

The ML velocity of the CoM differed in the presence of bilateral threats compared to the no threat, but not based on the direction of the threat. During the bilateral threat condition, participants reduced their ML CoM velocity compared to the ground-level condition, but this effect was only significant on the inner limb (*β* = 0.019, SE = 0.005, p < 0.001 for the inner limb; *β* = 0.01, SE = 0.006, p = 0.090 for the outer limb; Figure 8). There was no difference in ML CoM velocity between inner and outer threat conditions for either the inner or outer stance limbs (*β* = 0.005, SE = 0.005, p = 0.281 and *β* = 4.47e-04, SE = 0.006, p = 0.938, respectively).

**Figure 8.**
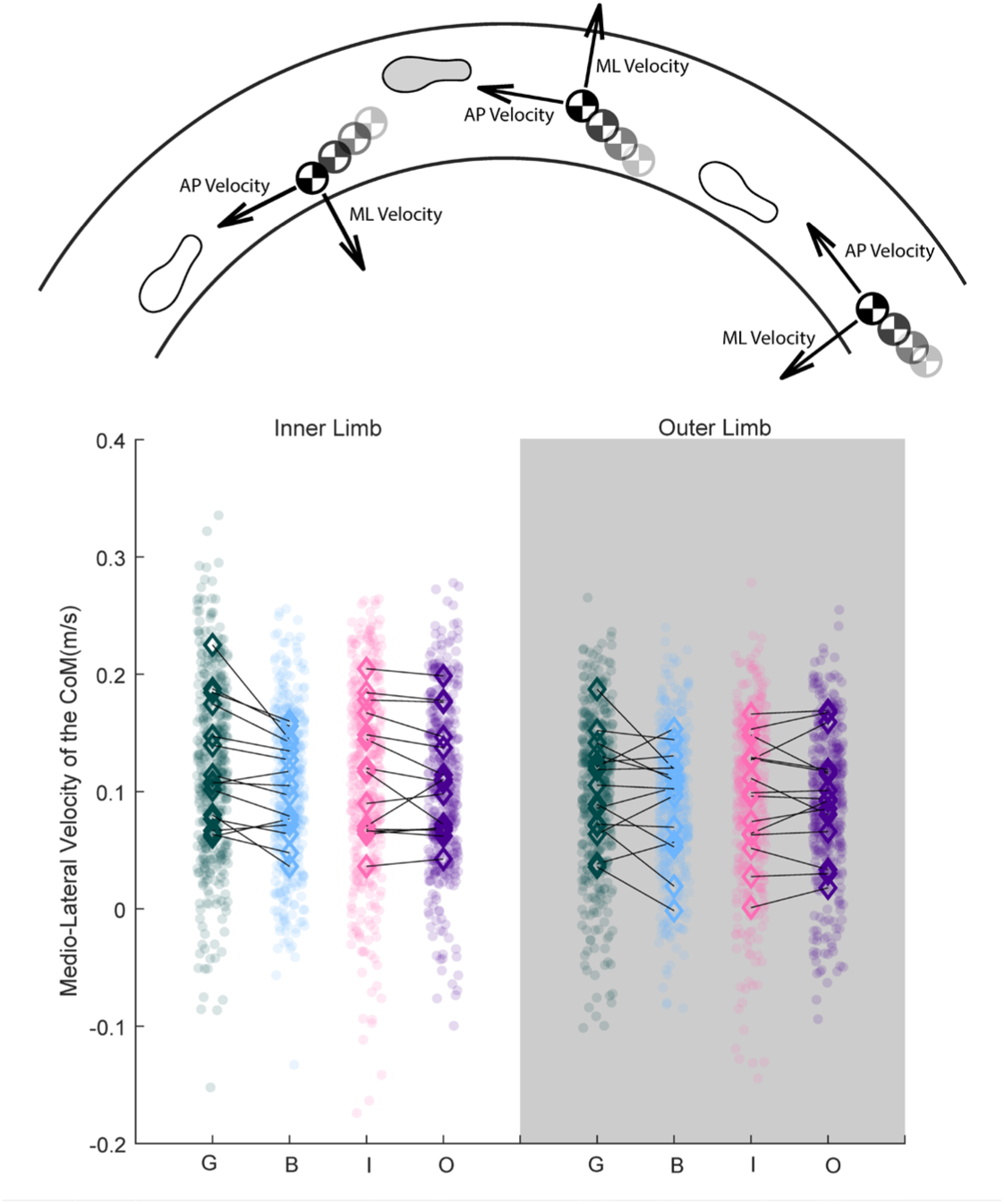
Mediolateral (ML) velocity of the center of mass (CoM) results for Experiment 2 during ground-level (G), bilateral threats (B), inner threat (I), and outer threat (O) for the inner (white shading) and outer limb (gray shading). ML CoM velocity was extracted at contralateral toe-off for each step. Data points represent individual steps across all participants, and the connected diamonds indicate within-participant mean values for each condition. Greater ML CoM velocities indicate lateral motion towards the edge ipsilateral edge of the walkway.

## DISCUSSION

We tested whether individuals alter their locomotor behavior to reduce the PoF when the perceived cost of falling increases. Based on the risk model and an assumption the individuals alter their behavior to maintain a constant level of risk, we expected participants to adopt compensatory strategies that minimized their probability of falling when walking in threatening environments with high perceived costs of falling. In Experiment 1, we observed that individuals altered their locomotor trajectory to move away from perceived threats that pose high costs (e.g., a high ledge), and that this change in trajectory incurred an implicit penalty (additional distance traveled orthogonal to the stated goal). In Experiment 2, we extended these results and observed that participants decreased the PoF in the direction of perceived threats. Furthermore, changes in the PoF were accomplished differently depending on the location of the threats. In unilateral threat conditions, participants adjusted their position in the walkway to move further away from the threat, like the results of Experiment 1, thereby increasing their MoF and decreasing the PoF. However, the bilateral threat condition did not afford such a solution. In this condition, participants moved their outer foot closer to the center of the walkway, took *narrower* steps, and decreased the ML CoM velocity when loading their inner limb. Taken together, our results support our hypothesis that individuals alter locomotor behavior in accordance with a risk model based on the probability of an action (i.e., the probability of falling off the edge of the walkway) and the perceived cost of that action (i.e., the perceived cost of falling off the edge of the walkway) [14, 36, 48].

We have provided evidence that humans adjust both their trajectory and balance control during walking to accommodate changes in the perceived cost of falling. Recent work in two-dimensional simulations has modeled compensatory adjustments to locomotor trajectories that scale with the magnitude of the threat [49]. Similarly, previous research on balance and locomotion in elevated physical or virtual environments has often emphasized postural strategies as a key compensatory mechanism to mitigate the increased threat of falling. For instance, individuals reduce CoM velocity in the direction of threats and take narrower steps when navigating height-related threats, aligning with our findings from Experiment 2 regarding bilateral threats condition [32, 50]. The present results suggest individuals alter locomotor trajectory and balance control in tandem.

While the present experiments were not designed to infer hierarchical control, results from Experiment 2 suggest that trajectory control may be a primary strategy to control one’s proximity to threats, based on differences in foot radial position and the lack of differences in ML CoM velocity during unilateral threat conditions. Results from the bilateral threat condition in Experiment 2 suggest that balance control mechanisms, specifically those that reduce the CoM velocity toward threats, are used as a secondary compensatory control strategy. Both compensations likely elicit additional (non-fall-related) penalties compared to ground-level walking. For instance, lateral stepping deviations in Experiment 1 resulted in longer walking distances, and changes in foot placement along the walkway in Experiment 2 necessitated longer walking distances, different curvatures, and different linear walking speeds. Increased walking distances may have also increased the metabolic and temporal demand of walking compared to ground-level conditions [36, 51]. Similarly, the compensations to balance control likely induced similar penalties to energetics and / or adaptability. During the bilateral condition in Experiment 2, a reduction in the ML CoM velocity was achieved by narrowing the base of support, indicated by a smaller distance between inner and outer foot radii. Since circumduction during swing is a contributor to the metabolic demand of walking, this narrow gait likely increased the energetic demand of walking[52]. Similarly, a reduction in ML CoM velocity could be considered as a “stiffening” strategy, where people restrict their CoM range of motion [31], resulting in a lower amplitude in postural sway. However, this stiffening strategy may not be beneficial in more complex tasks that require flexible, redundant locomotor control [12, 53]. In the future, researchers should explore how risk-based compensatory strategies change with larger competing implicit and explicit penalties (e.g., steeper metabolic or temporal penalties for changes in trajectory, perturbation and complex walking terrain for changes in balance control) to elucidate whether hierarchical control is used in different risk-based scenarios.

Speculative interpretations of past results when standing or walking with height-related threats have focused on the potential for individuals to freeze degrees of freedom to simplify the task when facing high height threat [54, 55]. Individuals under anxiety, potentially caused by height-related threats, could experience a disruption in an automated motor task and revert to “novice-like” motor behaviors by reducing degrees of freedom to simplify a motor task [11, 54, 55]. However, our results suggest an alternative explanation – participants reduce the CoM velocity not merely to simplify the task but specifically to reduce the probability of high-cost consequences (e.g., the probability of falling off a high cliff). This strategy of reducing degrees of freedom to alleviate the risk of falling complements existing theories and underscores the role of perceived risk in modulating locomotor strategies. We speculate that simplification and threat-avoidance are not mutually exclusive, and both may contribute to the observed behaviors.

Several factors need to be considered when interpreting the results. First, the use of a VR environment may not have elicited as convincing a perceived cost of falling as a physical environment would. Although our MRF-3 and RSME results indicate that both experiments successfully induced anxiety related to height (see Supplement), the high-height VR setting may not reflect risk perception in real-world environments. Second, we used only one height condition. Ideally, multiple height conditions would have been incorporated to explore whether locomotor adaptations scale with varying levels of perceived cost / threat as shown in mathematical simulations and standing postural control tasks [49, 56]. Consequently, while our findings align with predictions made by the risk model, we cannot draw definitive conclusions about the scaling of these changes across different levels of perceived threat (i.e., levels of perceived cost of falling). Such scaling could vary across the environmental threat or inter-individual weighting of threat (see Figure 1). We explored whether our results were influenced by visual height intolerance and found slightly larger effects in people with reported visual height intolerance compared to those without visual height intolerance (see Supplement), but future research should explore factors that may influence interindividual differences. Such factors may be targets for behavioral modification strategies, including mobility-enhancing interventions. Third, the relatively small sample size could limit the generalizability of our findings, particularly to other populations. However, this limitation is somewhat mitigated by replicating the results across different samples in related, but distinct experiments. Fourth, our results for the MoF and PoF in Experiment 2 relied on an approximation of the CoM using two positional markers, rather than a whole-body CoM. It is likely this approach biases our estimates of the MoF and PoF between participants. However, since all comparisons were assessing within-participant changes, it is unlikely that these errors influenced the conclusions. Additionally, the small magnitude of the PoF should be interpreted cautiously. As indicated in Figure 5, participants did not generally exhibit MoF values that were close to 0 (i.e., close to the edge of the walkway). Therefore, the PoF outcome was based on extrapolating the distribution of MoF to negative values, rather than direct observations of negative MoF values. Nonetheless, the results from the PoF complement the results of the MoF. In the future, researchers should address these limitations by incorporating a broader range of environmental conditions, larger sample sizes to further validate and extend our findings, and whole-body kinematic and kinetic data to completely characterize kinematic and kinetic compensatory strategies.

## CONCLUSIONS

This study reveals how individuals dynamically adjust their locomotor behavior to manage perceived fall risk, underscoring the significant role of perceived danger in shaping walking and balance strategies. Our findings demonstrate that in high-risk scenarios, people actively modify their trajectory and stability to mitigate changes in the perceived cost of falling, a behavior that strongly supports the principles of a risk-based balance control strategy. This research deepens our understanding of the interplay between risk perception and locomotor control and highlights human locomotor behavior in response to perceived threats. The implications of these findings may aid in the development of fall prevention strategies and mobility interventions that target an individual’s risk landscape, as well as robotic controllers that seek to mimic human locomotor decisions. By applying insights gained from this work, we can design more effective safety measures and environmental modifications that enhance mobility and reduce the risk of falls in various settings.

## ACKNOWLEDGEMENTS

We would like to express our gratitude to Tyler Ho for his contributions to figure design and to Benjamin Engel for assistance with the virtual reality environment.

## DATA, CODE AND MATERIALS

All data and analysis materials are available for Experiment 1 (doi: 10.6084/m9.figshare.27773973) and Experiment 2 (doi: 10.6084/m9.figshare.27774504).

