## Supplement for "Behavioral risk models explain locomotor and balance changes when walking at virtual heights"

### SUPPLEMENT A

Prior work has documented the effectiveness of the virtual reality environment used here as eliciting height-related anxiety [27, 32, 57]. To confirm the effectiveness of our virtual height manipulation for eliciting height-related anxiety, the Mental Readiness Form-3 (MRF-3) and Rating Scale of Mental Effort (RSME) were administered [45, 46]. In Experiment 1, the MRF-3 and RSME were administered after every walking pass in the first and third trial blocks (i.e., after every walk from one end of the walkway to the other). In Experiment 2, the MRF-3 and RSME were administered after every condition (i.e., after every 40 s walking trial). For both experiments, participants remained in the virtual environment while completing these questionnaires; the MRF-3 was administered verbally and the RSME was displayed in the virtual environment to ensure participants had consistent anchors for their responses.

Statistical comparisons were not completed. Descriptive statistics were interpreted for each experiment. Descriptively, participants reported greater anxiety and greater mental effort when walking at high height, compared to low height in Experiment 1 (**Table e1**).

**Table e1.** Comparison of MRF-3 and RSME Scores Across Low and High Height Conditions for Experiment 1 to Confirm the Fidelity of the Virtual Condition

| Statement | Low Height M (SD) | High Height M (SD) |
|---|---|---|
| "My thoughts were…" 0 (calm) - 10 (worried) | 1.8 (1.6) | 3.4 (2.4) |
| "My body feels…" 0 (relaxed) – 10 (tense) | 1.7 (1.6) | 3.7 (2.3) |
| "I am feeling…" 0 (confident) – 10 (not confident) | 0.8 (1.3) | 2.0 (1.8) |
| Mental effort allocated to task: 0 (none) – 150 (extreme) | 29.9 (27.7) | 45.0 (25.7) |

Similarly, increased anxiety and mental effort were reported in all three high height conditions (bilateral, inner, and outer threat conditions) compared to ground level conditions in Experiment 2 (**Table e2**). Participants reported the greatest anxiety and mental effort during the bilateral threat condition. Anxiety and mental effort were similar for the inner and outer threat conditions.

**Table e2.** Comparison of MRF-3 and RSME Scores Across Conditions Ground Level (G), Bilateral Threat (B), Inner Threat (I), and Outer Threat (O) to Confirm the Fidelity of the Virtual Condition

| Statement | G<br>M (SD) | B<br>M (SD) | I<br>M (SD) | O<br>M (SD) |
|---|---|---|---|---|
| “My thoughts were…” 0 (calm) - 10 (worried) | 1.3 (1.9) | 4.4 (2.7) | 2.9 (1.9) | 3.5 (2.4) |
| “My body feels…” 0 (relaxed) – 10 (tense) | 1.7 (1.8) | 4.8 (2.7) | 3.3 (2.2) | 3.7 (2.5) |
| “I am feeling…” 0 (confident) – 10 (not confident) | 1.3 (2.1) | 3.6 (2.0) | 2.7 (2.0) | 2.3 (2.2) |
| Mental effort allocated to task: 0 (none) – 150 (extreme) | 23.3 (13.6) | 50 (18.1) | 40.3 (20.1) | 42.7 (20.8) |

### SUPPLEMENT B

It is possible that some individuals would exhibit larger responses to the virtual height environment based on their own assessments of risk. For example, individuals with a particularly strong visual height intolerance (VHI, i.e., fear of heights), may have exhibited stronger responses to the environment and driven the results [58]. To explore whether the results presented in this study were significantly influenced by individual differences in VHI, we stratified participants in Experiment 2 based on their self-reported VHI [58]. A total of 8 people reported VHI (VHI+), and 7 people reported no VHI (VHI-). The statistical analysis described in the methods for the entire sample was completed separately for these stratified groups. To interpret the results, we qualitatively compared pair-wise effect sizes between the VHI+ and VHI- groups.

Overall, effects between the bilateral and ground conditions were similar in direction, but smaller in magnitude, in the VHI- participants (Table e3) compared to the VHI+ participants (Table e4). Qualitatively similar results in both magnitude and direction were observed between VHI+ and VHI- participants for effects between inner and outer threat conditions.

**Table e3.** Descriptive means (M) and standard deviations (SD) and pair-wise effect sizes for margin of falling (MoF), probability of falling (PoF), radial foot placement, and mediolateral (ML) center of mass (CoM) velocity, across different threat conditions and stance limb for participants without visual height intolerance (VHI-).

| | Stance Limb / Edge | Ground Level M (SD) | Bilateral Threat M (SD) | Hedge's *g* ES | Inner Threat M (SD) | Outer Threat M (SD) | Hedge's *g* ES |
|---|---|---|---|---|---|---|---|
| Margin of Falling (MoF) (m) | Inner | 0.14 (0.03) | 0.12 (0.02) | 0.39 | 0.16 (0.04) | 0.10 (0.02) | 2.23 |
| | Outer | 0.23 (0.02) | 0.21 (0.03) | 0.58 | 0.18 (0.03) | 0.26 (0.02) | 2.48 |
| Probability of Falling (PoF) (%) | Inner | 8.49e-03 (1.72e-02) | 6.14e-05 (1.49e-04) | 0.93 | 4.71e-04 (1.22e-03) | 7.86e-02 (1.05e-01) | 1.81 |
| | Outer | 2.17e-08 (3.79e-08) | 8.72e-15 (8.39e-15) | 1.65 | 3.57e-05 (6.98e-05) | 8.70e-04 (2.30e-03) | 0.64 |
| Radial Foot Placement (m) | Inner | 1.35 (0.05) | 1.34 (0.03) | 0.26 | 1.38 (0.05) | 1.31 (0.02) | 1.63 |
| | Outer | 1.45 (0.03) | 1.42 (0.03) | 0.69 | 1.46 (0.04) | 1.41(0.04) | 1.78 |
| ML CoM Velocity (m/s) | Inner | 0.11 (0.03) | 0.11 (0.03) | 0.28 | 0.11 (0.04) | 0.11 (0.03) | 0.09 |
| | Outer | 0.09 (0.03) | 0.09 (0.04) | 0.01 | 0.09 (0.04) | 0.08 (0.04) | 0.21 |

Margin of Falling (MoF): Represents the radial distance from the edge of the walkway to the extrapolated Center of Mass (XcoM) at contralateral toe-off, indicating the stability margin in each condition.
Probability of Falling (PoF): The estimated probability, expressed as a percentage, that a participant will fall under each condition.
Medial-lateral velocity of the Center of Mass (ML CoM Velocity): The medial-lateral velocity of the Center of Mass (CoM), measured in meters per second (m/s).

**Table e4.** Descriptive means (M) and standard deviations (SD) and pair-wise effect sizes for margin of falling (MoF), probability of falling (PoF), radial foot placement, and mediolateral (ML) center of mass (CoM) velocity, across different threat conditions and stance limb for participants with visual height intolerance (VHI+).

| | Stance Limb / Edge | Ground Level M (SD) | Bilateral Threat M (SD) | Hedge's *g* ES | Inner Threat M (SD) | Outer Threat M (SD) | Hedge's *g* ES |
|---|---|---|---|---|---|---|---|
| Margin of Falling (MoF) (m) | Inner | 0.15 (0.03) | 0.13 (0.02) | 0.90 | 0.18 (0.02) | 0.12 (0.02) | 2.59 |
| | Outer | 0.19 (0.04) | 0.22 (0.03) | 0.97 | 0.17 (0.03) | 0.23 (0.03) | 1.65 |
| Probability of Falling (PoF) (%) | Inner | 1.24e-03 (3.49e-03) | 1.70e-04 (4.47e-04) | 0.36 | 4.38e-05 (9.49e-05) | 5.01e-02 (1.41e-01) | 0.62 |
| | Outer | 1.43e-05 (3.59e-05) | 7.84e-14 (2.02e-13) | 3.12 | 1.14e-04 (2.95e-04) | 1.81e-04 (5.10e-04) | 1.01 |
| Radial Foot Placement (m) | Inner | 1.37 (0.02) | 1.37 (0.03) | 0.16 | 1.40 (0.02) | 1.34 (0.03) | 2.01 |
| | Outer | 1.45 (0.03) | 1.42 (0.03) | 1.07 | 1.47 (0.04) | 1.41 (0.04) | 1.49 |
| ML CoM Velocity (m/s) | Inner | 0.13 (0.06) | 0.10 (0.05) | 0.42 | 0.12 (0.06) | 0.11 (0.06) | 0.09 |
| | Outer | 0.11 (0.05) | 0.09 (0.05) | 0.28 | 0.10 (0.05) | 0.11 (0.05) | 0.15 |

Margin of Falling (MoF): Represents the radial distance from the edge of the walkway to the extrapolated Center of Mass (XcoM) at contralateral toe-off, indicating the stability margin in each condition.
Probability of Falling (PoF): The estimated probability, expressed as a percentage, that a participant will fall under each condition.
Medial-lateral velocity of the Center of Mass (ML CoM Velocity): The medial-lateral velocity of the Center of Mass (CoM), measured in meters per second (m/s).
